# Variation in tolerance to parasites in natural Asian tiger mosquito populations and its effect on vectorial capacity

**DOI:** 10.1101/445759

**Authors:** Guha Dharmarajan, Kathryne D. Walker, Tovi Lehmann

## Abstract

The vectorial capacity of mosquitoes depends upon the magnitude of reduction of parasite load upon infection through resistance mechanisms (e.g., immune-mediated killing) and the ability of mosquitoes to offset infection-mediated costs through tolerance mechanisms (e.g., tissue repair). Here we use a common-garden experimental framework to measure variation in resistance and tolerance to dog heartworm (*Dirofilaria immitis*) between natural *Aedes albopictus* mosquito populations representing areas of low and high transmission intensity. Our data revealed that survival to the extrinsic incubation period, the earliest time point at which infective L3 larvae develop, significantly differed between populations (ranging from 10-60%) when mosquitoes infected with *D. immitis* at both the low (15 microfilaria/μl blood) and high (30 microfilaria/μl blood) infection dose (Dose: χ^2^ = 191.473; *P* < 0.001; Population: χ^2^ = 24.485; *P* = 0.001; Dose × Population: χ^2^ = 35.566; *P* = 0.001). Contrary to expectations, we found that mosquito populations with highest resistance (i.e., greatest reduction in parasite load) also exhibited highest mortality upon infection (*F*_1,12_ = 6.781, *P* = 0.023; Dose: *F*_1,12_ = 6.747; *P* = 0.023; Mortality × Dose: *F*_1,12_ = 0.111, *P* = 0.744). Expressing the effect of the number of killed (N_KILLED_) and live (N_LIVE_) parasite on survival of mosquitoes from the different population, we document a significant inter-population variation in the survival cost of additional parasite (i.e., tolerance to infection (N_LIVE_ × Population: χ^2^ = 22.845; *P* = 0.002; N_KILLED_ × Population: χ^2^ = 31.959; *P* = < 0.001; N_LIVE_ × N_KILLED_ × Population: χ^2^ = 22.266; *P* = 0.002), in conjunction with negative relationship between tolerance and resistance (Resistance: *F*_1,12_ = 11.870, P = 0.005; Dose: *F*_1,12_ = 16.0170, P = 0.002; Resistance × Dose: *F*_1,12_ = 9.699, P =0.009). Importantly, populations from areas with high transmission intensity (as measured by parasite prevalence in dogs) showed elevated tolerance (Prevalence: *F*_1,12_ = 9.5, P = 0.012; Prevalence^2^: *F*_1,12_ = 4.353, P = 0.064; Dose: *F*_1,12_ = 38.855, P = <0.001), and these populations were also associated with increased vectorial capacity (Tolerance: *F*_1,12_ = 8.175, *P* = 0.014; Dose: *F*_1,12_ = 0.005, *P* = 0.946; Tolerance × Dose: *F*_1,12_ = 0.920, *P* = 0.356). Consequently, our data indicate that spatial variation in disease transmission intensity is linked to the evolution of tolerance in natural mosquito populations, which in turn can feedback to impact disease risk.

## Introduction

Animal defense against parasites has been typically equated with host resistance (i.e., mechanisms that directly reduce parasite burden), but host tolerance, which has similar benefit to the host by minimizing harm (fitness cost) inflicted by the parasites (e.g. tissue repair) has been mostly ignored^1–5^. The importance of host tolerance in plants has long been recognized^6^, but animal ecologists have only recently started investigating the role tolerance plays in shaping host-parasite interactions (see references above). The defense responses of mosquito and other arthropod vectors against parasites has long been recognized central to understanding disease transmission and for the development of novel disease control strategies. Previous research has focused on resistance of mosquito vectors by immune and other defense (eg., cirabrial armature, peritrophic membrane) mechanisms that reduce the number/load of developing parasites^7,8^. While the role of tolerance as an alternative pathway for the vector to cope with parasite mediated damage has been mentioned^9^, to our knowledge, there have been no studies to evaluate tolerance in mosquitoes and its role in disease transmission‥ Both resistance and tolerance can act to improve the fitness of infected mosquitoes, yet these strategies have distinct evolutionary dynamics due to their differential effects on parasite fitness^1–4,10^. Because resistance negatively affects parasite fitness this strategy is expected to lead to antagonistic co-evolution between the vector and parasite (Red Queen dynamics^11^). Alternatively, by reducing parasite-mediated fitness costs rather than parasite load, tolerance is expected to have a neutral (or even positive) effect on parasite fitness^11^. Free from antagonistic co-evolution of the parasite, theory predicts a rapid evolution of tolerance against parasites, especially as the risk of infection increases^12^. To evaluate the roles that resistance and tolerance play in shaping mosquito-filaria relationships we estimated variation in these traits among eight natural populations of the Asian tiger mosquito, *Aedes albopictus* (Fig. 1A), infected with the dog heartworm (*Dirofilaria immitis*).

**Fig. 1.**
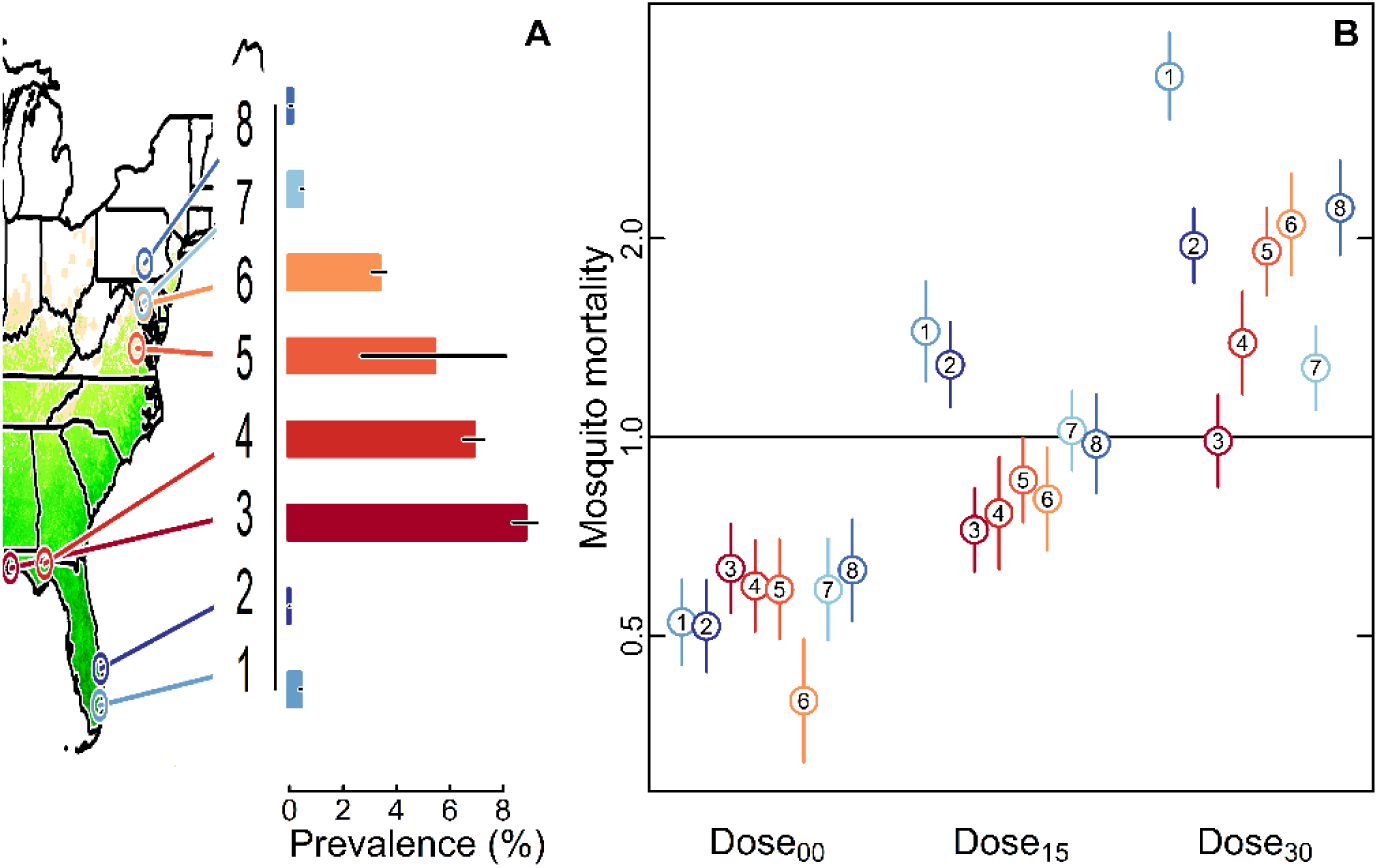
*Aedes albopictus* populations and their mortality patterns when infected with *Dirofilaria immitis*. **(A)** Map of locations from which the eight populations were sampled (left) with background depicting the range distribution of *A. albopictus* [27]. The bar graph shows the county-level prevalence of *D. immitis* antibodies in dogs (N = 19,570). **(B)** Population differences in daily mortality hazards among uninfected mosquitoes (Dose_00_) and mosquitoes infected using 15 and 30 *D. immitis* microfilaria/μl of blood (Dose_15_ and Dose_30_, respectively) compared to the hazard across all populations and doses (horizontal line). Mosquitoes at Dose_00_ and Dose_30_ had the lowest mortality (~2 fold decrease from baseline hazard) and highest (~2 fold greater than baseline hazard) mortality, respectively (Table S3). Error bars are standard errors of the mean and population numbers follow Fig. 1A.

*Aedes albopictus*, one of the most important disease vectors globally, is implicated in the transmission of *D. immitis* in dogs as well as Chikungunya, West Nile, Dengue, and Zika viruses in humans^13,14^. The *Aedes-Dirofilaria* system was chosen for this study because it is a natural vector parasite system^13^ wherein the parasite exerts high fitness costs on its vector: a modest exposure to 15 *D. immitis* microfilaria (mf)/μl of blood leads to ~60% mortality in *A. albopictus* prior to mosquito reproduction (i.e., within three days post-infection^15^). Additionally, the risk of mosquito exposure to *D. immitis* markedly varies amongst populations as measured by the spatial heterogeneity in prevalence of *D. immitis* in dogs^16,17^, providing a “natural experiment” of the role of parasite selection intensity on the evolution of tolerance and resistance. Specifically, we addressed the following questions: Do these populations differ in their survival to infection and how that variation relates to parasite load? If survival depends on resistance and tolerance how does that variation relate to the mosquito risk of exposure to the parasite? Finally, what are the implications of these answers for the mosquito vectorial capacity?

## Results and Discussion

Employing a common garden experimental design, we measured mosquito resistance and tolerance in F2 offspring of eight mosquito populations experimentally infected with *D. immitis* (Fig. 1A; Fig. S1). These mosquito populations were selected to span areas representing low to high risk of parasite exposure, as determined by prevalence of *D. immitis* in dogs (0-9%, Fig. 1A; Table S1; (18)). Blood feeding and experimental infection of F2 offspring from each population was carried out via membrane feeding using three concentrations of *D. immitis* mf: 0, 15 and 30 mf/μl (Table S2). Infected mosquitoes revealed minimal between-population (and between-cage) variation in the exposure to parasites (i.e., microfilaria; mf) as measured by the zero-hour mf counts within dose (Dose > 0; Replicate: *F*_1,169_ = 177.65; *P* = 0.501, Population: *F*_7,169_ = 2469.88; *P* = 0.506; Population × Dose: *F*_7,169_ = 2231.80; *P* = 0.576; Fig. S2). Mosquitoes exposed to low vs. high dose (15 vs. 30 mf/μl) had significantly different initial infection intensities (*F*_1,169_ = 24373.23; *P* <0.001; Fig. S2).

### Mosquito mortality

Consistent with previous studies^15^, mosquito mortality dramatically increased by infection, showing a highly significant effect of infection dose (Fig 1B; Fig. S3; Table S3; χ^2^ = 191.473; df = 2; *P* < 0.001). Notably, there was also a strong interactive effect of dose and population on mosquito mortality (Fig. 1B; Fig. S3; Table S3; Population: χ^2^ = 24.485; df = 7; *P* = 0.001; Dose × Population: χ^2^ = 35.566; df = 14; *P* = 0.001), indicating infection with *D. immitis* does not affect fitness uniformly across different mosquito populations (Fig 1B).

### Resistance

The conventional view predicts that resistance would improve mosquito fitness in the face of parasite infection. Thus, we tested if mosquito populations differed in the magnitude of resistance, measured as the magnitude of reduction in parasite load at the extrinsic incubation period (EIP; i.e., the earliest time point at which infective L3 larvae develop) compared to the initial (time zero) parasite exposure for a given population and dose (see Materials and Methods). Mosquito populations varied significantly the rate at which they killed parasites (Fig. 2A; Fig. S4; Table S4; Day × Population: χ^2^ = 18.820; df = 7; *P* = 0.009). Thus, despite being infected by a similar number of mf at time zero, the populations differed in parasite load at the EIP (Fig 2A). Surprisingly, however, mosquito populations with highest resistance (i.e., greatest reduction in parasite load) also exhibited highest mortality upon infection; Fig 2B and C; Mortality: *F*_1,12_ = 6.781, *P* = 0.023; Dose: *F*_1,12_ = 6.747; *P* = 0.023; Mortality × Dose: *F*_1,12_ = 0.111, *P* = 0.744). This pattern was not driven by selection (i.e., non-random mortality due to infection burden) given that parasite loads did not differ between dead mosquitoes and those that were live but censored during the course of the experiment (Fig. S5; *F*_2,409_ = 0.815; *P* = 0.443).

**Fig. 2.**
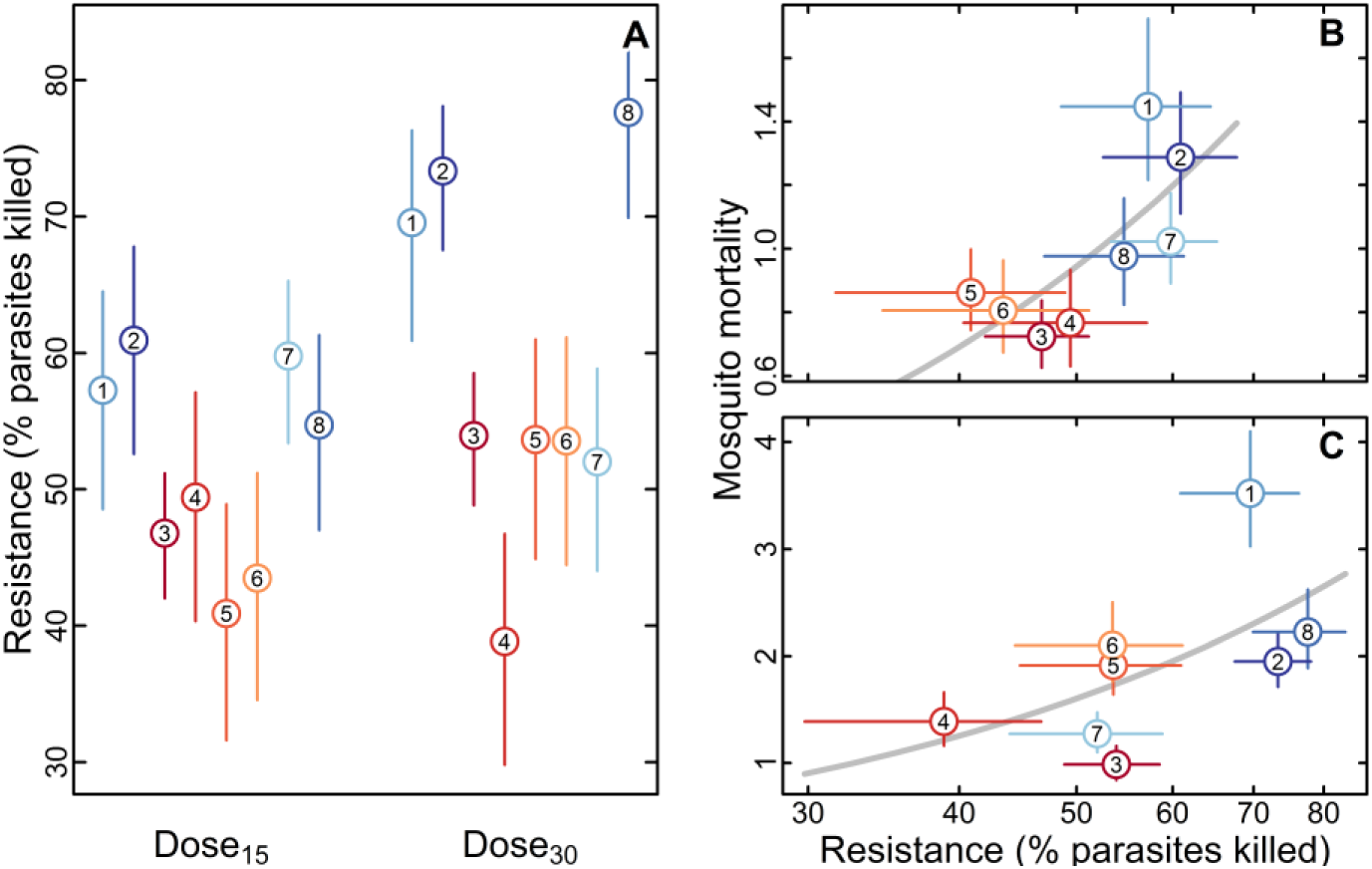
Resistance to *Dirofilaria immitis* infection amongst the eight *Aedes albopictus* populations. **(A)** Population differences in resistance measured as the magnitude of reduction in parasite load at the extrinsic incubation period compared to the initial (time zero) infection load (Fig. S2) in mosquitoes infected using 15 and 30 *D. immitis* microfilaria/μl of blood (Dose_15_ and Dose_30_, respectively) (Table S4). **(B** and **C)** Relationship between resistance and mortality hazard in mosquitoes infected with 15 (B) and 30 (C) microfilaria/μl of blood. Regression model predictions are also shown (gray line). Error bars are standard errors of the mean and population numbers follow Fig. 1A.

### Tolerance

The positive relationship between resistance and mortality could be driven by differences in the ability of mosquitoes to heal tissue damage caused by the parasites or the “collateral damage” that could result from activation of anti-parasite immunity^10,18,19^. To determine the presence of a tolerance response and assess its contribution to infected mosquito survival, we estimated the population-specific effects of killed-and live-parasites on mortality, such that tolerance was evident in the presence of a significant between-population variance in the effect of live parasites on mortality, after accommodating the effect of killed parasites (i.e., resistance) (see Materials and Methods). Thus, populations that have evolved higher tolerance were expected to exhibit significantly lower effects of live parasites on mortality (i.e., distinct slope and/or intercept). A systematic change in this effect among populations, with respect to the risk of exposure to the parasite, may further demonstrate evolution of tolerance and suggest an evolutionary process that shapes it. Mosquito mortality was strongly affected by both the number of live and killed parasites. (Table S5; N_LIVE_ × Population: χ^2^ = 22.845; df = 7; *P* = 0.002; N_KILLED_ × Population: χ^2^ = 31.959; df = 7; *P* = < 0.001; N_LIVE_ × N_KILLED_ × Population: χ^2^ = 22.266; df = 7; *P* = 0.002). In the case of macroparasites, like *D. immitis*, which do not reproduce in the vector, at a given infection dose, an increase in the number of live parasites at a specific time point is necessarily associated with a reduction in the number of killed parasites (i.e., a reduction in resistance).

To elucidate how parasite load affected mosquito mortality, we compared the effect of a unit increase in live parasite and a unit decrease in killed parasites when mosquitoes are exposed to low (15 mf/μl) or high (30 mf/μl) infection dose. Our results indicated that mosquito populations differed in levels of tolerance. These differences were especially pronounced when comparing the high exposure populations (i.e., Pop 3, 4, 5 and 6) with the low exposure populations (i.e., Pop 1, 2, 7, 8). For example, considering Pop 3 infected at the low infection dose (i.e., 15 mf/μl), this population has a higher tolerance to infection compared to Pop 1, 2 and 8 (Fig. 3 A,B and D), but not Pop 7 (Fig 3 C), with consistently lower mortality at all levels of live parasites, except in situations when mosquitoes kill over 80% of their parasites (a level of resistance higher than maximum resistance in any population; Fig.2A). These patterns were qualitatively similar when comparing the other high exposure populations (i.e., Pop 4, 5 and 6) infected at the low infection dose (Fig. S6-8). At the high infection dose, we found greater variability in tolerance. For Pop 3, these patterns were similar to the low infection dose (Fig. 1 E-H), but for Pop 5 that wassimilar to Pop 3 at the low infection dose, no differernce was found in comparisons with the four low exposure populations at the high infection dose (Fig. S7), indicating that resistance is equivalent to tolerance at high infection dose in terms of survival. A probable tradeoff between resistance and tolerance is indicated by a significant negative relationship between resistance and tolerance across all populations (Fig. S9B and C). Interestingly, there was also a significant positive, albeit non-linear, relationship between observed prevalence of *D. immitis* in dogs and the observed levels of tolerance, especially at the low infection dose (Prevalence: F = 9.5, P = 0.012; Prevalence^2^: F = 4.353, P = 0.064; Dose: F = 38.855, P <0.001; Fig. S9 D and E).

**Fig. 3.**
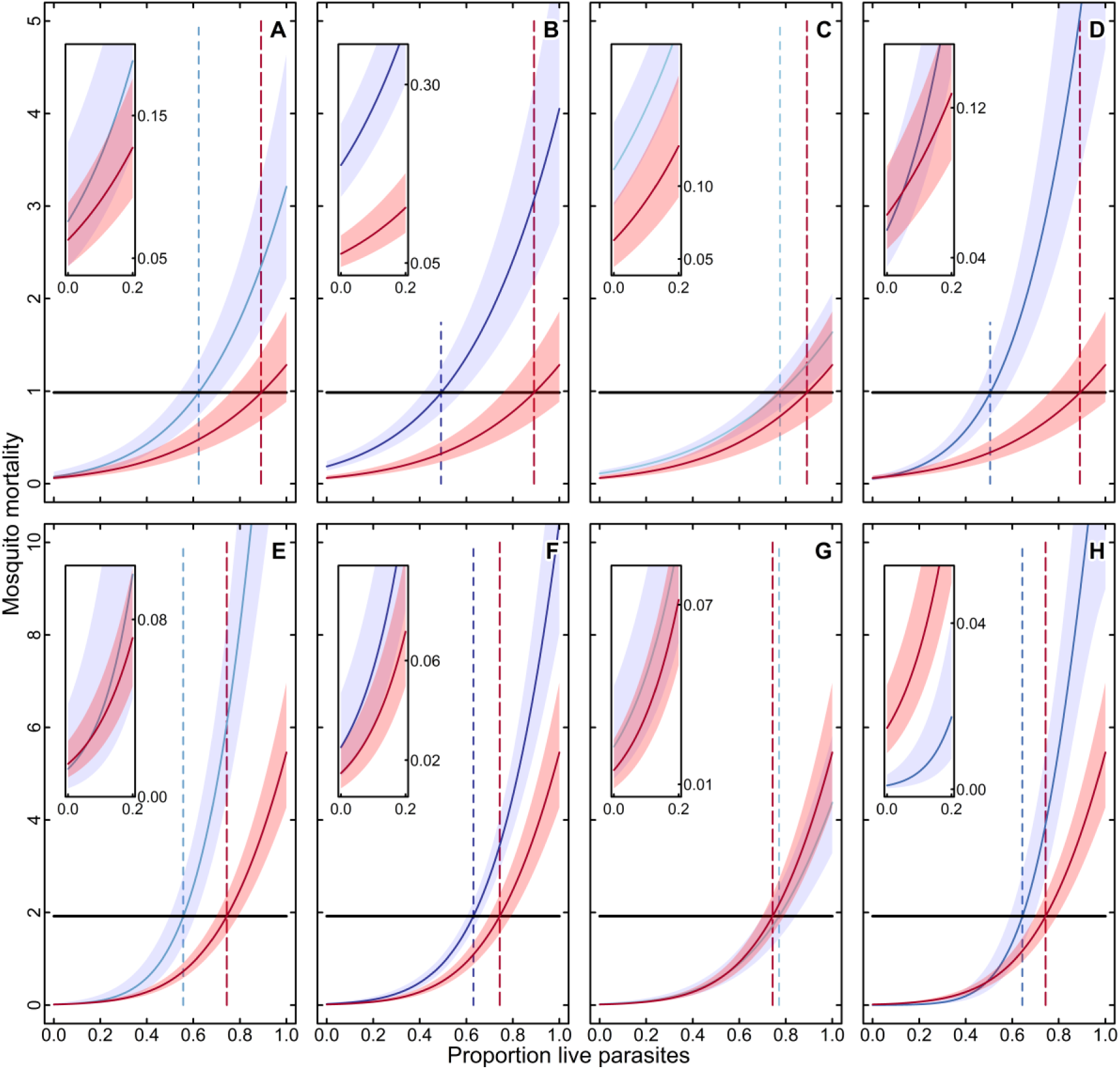
Tolerance to *Dirofilaria immitis* infection demonstrated as between-population variation in fitness of *Aedes albopictus* infected with the same load of live parasites, accommodating their resistance as predicted using Cox-Proportional Hazards Mixed Effects models (see text). Differences in mortality hazard as the proportion of live parasites increases for mosquitoes infected at two *D. immitis* infection doses: 15 (**A-D**) and 30 (**E-H**) microfilaria/μl of blood (Table S5). The graphs compare patterns of mortality between one high exposure population (Pop 3; red symbols) and the four low exposure populations (blue symbols): Pop 1 (**A** and **E**), Pop 2 (**B** and **F**), Pop 7 (**C** and **G**) and Pop 8 (**D** and **H**). Error bands are standard errors of the mean and population numbers follow Fig. 1A. Also represented are the maximum live parasite loads (dashed vertical lines) at which the population-specific mortality hazard is below average mortality hazard across all populations for a given infection dose (horizontal lines). The insets show mortality curve details at the highest levels of resistance (i.e., proportion of live parasites ≤ 0.2). Comparisons of the three other high exposure populations (i.e., Pop 4, 5, and 6) with the four low exposure populations are shown in Fig. SA-SC.

### Vectorial capacity

The differences between populations in terms of tolerance (and resistance) must have epidemiological implications in terms of disease transmission intensity due to variation in vectorial capacity, especially because levels of resistance (i.e., the reduction in parasite load; Fig. 2A) and tolerance (i.e., the relative survival at peak infection load compared to the baseline survival; Fig. S9A) were negatively correlated (Fig. S9B and C). We preferentially use the term vectorial capacity over vector competence, since the former incorporates the longevity of the vector on parasite transmission risk while the latter only refers to the ability of the vector to support parasite development to the infectious stage^20^. Vectorial capacity, and thus force of parasite transmission, is affected by the probability of a mosquito surviving to EIP and the number of infective L3 larvae in the surviving mosquitoes^21^. We thus estimated vectorial capacity as the joint probability of a mosquito surviving to EIP and of mf developing to infective L3 larvae in the survivors (see Materials and Methods). We found that vectorial capacity was impacted by both infection dose and population (Fig. 4A), a pattern driven by the effects of these variables on probability of survival to EIP (Fig. S10A; Table S6; Dose: χ^2^ = 45.188; df = 1; P <0.001; Population: χ^2^ = 25.462; df = 7; P = 0.001) and development of mf to L3 (Fig. S10B; Table S7; Dose: χ^2^ = 5.569; df = 1; P = 0.018; Population: χ^2^ = 17.455; df = 7; P = 0.015). As predicted, tolerance was positively associated with vectorial capacity, irrespective of dose (Fig. 4B and C; Tolerance: *F*_1,12_ = 8.175, *P* = 0.014; Dose: *F*_1,12_ = 0.005, *P* = 0.946; Tolerance × Dose: *F*_1,12_ = 0.920, *P* = 0.356). It is worth noting that mosquitoes surviving to EIP had no infective larvae in two populations (1 and 2; Fig. 4A). While none of the individuals in these two populations harbored infective larvae, most retained earlier stages (e.g., L2s) of infection, and individuals from these populations did harbor infective L3 at the lower infection dose (Fig. 4A). Given that less than 10% of mosquitoes in these populations (i.e., 1 and 2) eliminated all their parasites at the high infection dose (Table S2), the low vectorial capacity is not due to complete resistance to *D. immitis* but rather to higher mortality while infected, underlining the importance of tolerance in understanding vector-borne disease dynamics.

**Fig. 4.**
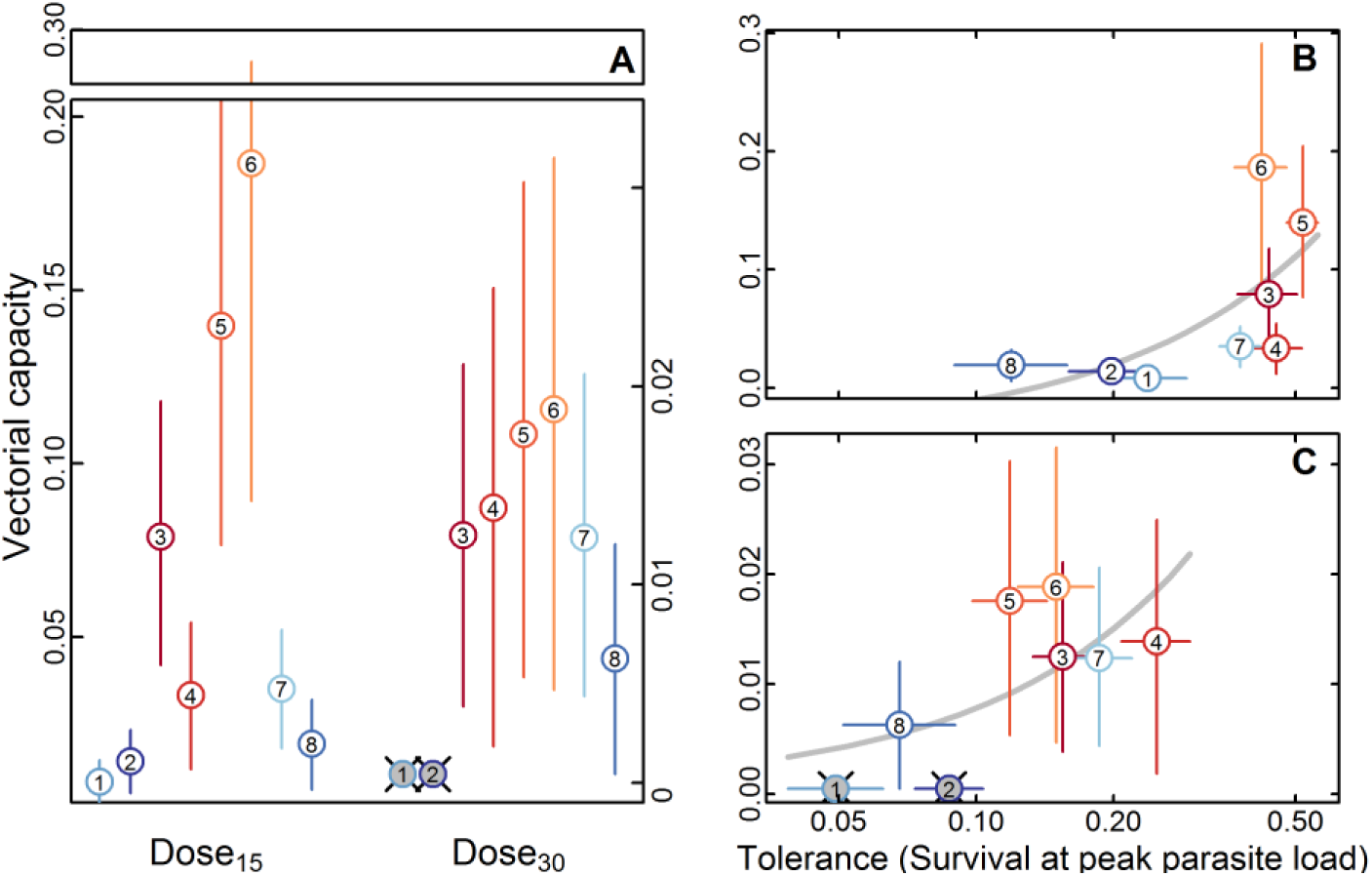
*Dirofilaria immitis* vectorial capacity of the eight *Aedes albopictus* populations. **(A)** Population differences in vectorial capacity measured as the probability of a microfilaria developing to L3 given the probability of mosquito survival to the extrinsic incubation period in mosquitoes infected using 15 and 30 *D. immitis* microfilaria/μl of blood (Dose_15_ and Dose_30_, respectively). Note that at Dose_30_, no infective L3 larvae were recovered from mosquitoes surviving to EIP, and these populations were dropped from the analyses (gray symbols) (Table S6 and S7). **(B** and **C)** Relationship between parasite tolerance, measured as the relative wait time between mortality events at peak infection load, and vectorial capacity in mosquitoes infected with 15 (B) and 30 (C) microfilaria/μl of blood. Regression model predictions are also shown (gray line). Error bars are standard errors of the mean and population numbers follow Fig. 1A.

## Conclusion

Mosquitoes constitute the most important group of disease vectors transmitting numerous diseases of global public health importance, such as malaria and lymphatic filariasis, as well as arboviruses such as Chikungunya, Dengue, Japanese encephalitis and Zika viruses^22–24^. However, this disease burden is not distributed uniformly at either global or local scales because mosquito-borne disease risk can be affected by spatial variation in many environmental and socioeconomic factors^16,17,25^. While such spatial heterogeneities in infection risk have obvious implications from a public health perspective, their effects on the vectors remain unclear.

Our results reveal that a higher risk of exposure to *D. immitis* predicted lower mortality under infection due to high parasite tolerance and low resistance (Fig S9 D and E). The divergent effects of resistance and tolerance on parasite fitness is particularly important in vector-borne disease dynamics because former strategy is expected to reduce vectorial capacity (due to a reduction of parasite burden), while the latter is expected to increase it (due to increased survival despite high parasite burdens)^9^. Consequently, parasite tolerance in mosquitoes has important implications for public health^10,26^, not only because vector-borne disease transmission and control hinge on vectorial capacity, which is shaped by the balance between vector resistance and tolerance, but also because even slight increases in transmission efficiency can enable establishment of novel parasites^27^. Additionally, because of the absence of counter evolution by parasites^1–4,10^, tolerance is expected to evolve more readily than resistance as might be the case here. The observed variation in resistance and tolerance could have evolved under different scenarios. It is tempting to propose that a major driver of these patterns is the spatial variation in risk of infection with the highly pathogenic *D. immitis*, because infection with *D. immitis* is likely to act as a potent selection pressure on *A. albopictus*, due to the high mortality in infected mosquitoes^15^ (Fig. 1A). Additionally, *A. albopictus* is naturally infected with *D. immitis*, with infection rates reported to range between 0-2%^13^, though the high mortality in infected mosquitoes suggests that these rates may greatly underestimate actual mosquito exposure. However, we cannot rule out selection by another parasite (affecting the aquatic or the adult stages) whose prevalence covaries with that of *D. immitis*, or even the mediation of non-parasitic agents. Consequently, additional studies are required to test the causal link between exposure risk to *D. immitis* and the evolution of tolerance in *A. albopictus* as described here.

Pioneering studies have revealed that variation in tolerance in other insects (e.g., *Drosophila melanogaster*^28^). However, the ability of mosquitoes to ameliorate the negative fitness consequences of infection through tolerance mechanisms has been virtually ignored (but see *REFS* 3, 4). This study provides evidence of the importance of tolerance for vector-pathogen interactions in natural mosquito populations. Given that *A. albopictus* has been introduced from Asia into the United States only recently (i.e., around 1985^31^) it is conceivable that the evolution of resistance and tolerance to infection is ongoing. However, since the experiments in this study were undertaken with F2 offspring under “common garden” conditions, the differences in phenotype reflect underlying genetic differences among the populations. Thus, our data indicate that differences in parasite selection pressure can lead to rapid divergence in the evolution of anti-parasite defense strategies in mosquitoes, highlighting the continued importance of field-based ecological and evolutionary studies in vector-parasite systems^32^.

The correlation between the degree of tolerance and the intensity of transmission (measured independently as the prevalence of *D. immitis* in dogs) suggests a role for tolerance for generating the spatial heterogeneity of dog heartworm, due to positive feedback between risk of acquiring the infection and vectorial capacity. Whether the parasite evolves to promote tolerance in the vector remains a mystery, but because tolerance benefits both vector and parasite, understanding the role of the parasite in the evolution of tolerance is a promising new frontier to improve our understanding of vector-borne disease dynamics in natural populations.

## Materials and Methods

### Collection and maintenance of mosquitoes

All experiments were carried out using eight *A. albopictus* lines specifically collected for this study from natural populations between July and September 2011 (see below). The Liverpool Blackeye strain of *A. aegypti* was used as a positive control^33^. The eight *A. albopictus* lines were established from mosquitoes collected from sites (Fig. 1A), representing a broad spectrum of risk of infection of *A. albopictus* with *D. immitis*, based on seroprevalence of *D. immitis* antibodies in dogs at the regional and county levels (Fig. 1A; Table S1). Each line was established from a collection of over 300 wild adult female mosquitoes. All field caught individuals were blood fed on chickens, and a random subset of 75-100 fully engorged females were aspirated into individual 50 ml tubes lined with paper towels for oviposition. Three days post feeding (dpf) we added 10 ml of water to each tube and mosquitoes were allowed to oviposit for 3 days. After oviposition species identity was confirmed using standard keys^34^. We randomly selected 50 *A. albopictus* females that oviposited > 25 eggs (F1 generation) and their eggs were used to establish the laboratory lines for each site. Thereafter, the lab lines were maintained using a large number of breeders (>1000 parents/generation) to minimize loss of genetic diversity due to genetic drift, and experiments were performed using eggs from the F1 generation adults. All larval rearing and adult maintenance were undertaken using standard insectary protocols and environmental conditions (27 °C; 75% humidity; 12:12 L:D diurnal cycle).

### Membrane feeding and infection of mosquitoes

Eggs from F1 generation adults from each line were synchronously hatched under standard conditions to produce F2 adults. These adults were maintained in one-gallon plastic containers with mesh tops at a density of *ca* 200 individuals/gallon with *ad libitum* access to sugar pads (10% Karo syrup) for 7 days prior to the experiment. One day prior to membrane feeding 60 female mosquitoes were transferred to ~500 ml plastic containers with mesh tops (henceforth “cages”). Twelve hours prior to membrane feeding, the sugar-pads on each cage were replaced by pads soaked in distilled water, and these water-soaked pads were removed six hours prior to membrane feeding. Mosquitoes in each cage were allowed to feed for 30 minutes on a hog-gut membrane stretched over an inverted water-jacketed glass feeder maintained at 40 °C. Each feeder was filled with 250 μL dog blood containing 1 mM ATP (as a phagostimulant^35^). Preparation of the dog blood for membrane feeding is described below. Briefly, we obtained *D. immitis* infected and uninfected dog blood from the Filariasis Research Reagent Resource Center, Athens, Georgia. To control for potential differences in infected and uninfected blood (e.g. nutritional differences) used in our experiment we isolated the microfilaria (mf) from the infected dog blood with minimal cell debris by membrane filtration using standard protocols^36^, and reconstituted in 2 ml of uninfected dog blood (this blood is henceforth designated “mf-blood”). The concentration of mf was determined as the average of 10 counts of 2 μL aliquots under 100X magnification. The treatment groups were fed a mixture of mf-blood and uninfected blood (proportions being diluted to the required mf dose; see below), and control (uninfected) groups were fed pure uninfected blood. Using this protocol we ensured that the treatment group and control group fed on the same blood, the only difference being that the blood fed to the treatment group contained a pre-determined concentration of mf.

### Experimental procedures

Our experiment was primarily designed to test if infection with *D. immitis* differentially affected survival among mosquito populations due to differences in resistance and tolerance. The experiment consisted of 2 replicates; each replicate consisted of three mf Doses (i.e., 0, 15 and 30 mf/μl). These infection doses are within the range of microfilaremia observed in naturally infected dogs^37–40^. In each replicated treatment, we fed 60 mosquitoes from each line in separate cages (see above). Approximately two hours after feeding we removed all unfed individuals and 6 fed individuals/cage to estimate the average number mf that mosquitoes in each cage were exposed to (in 3 cages <24 individuals fed and only 4 individuals were removed). These mosquitoes were stored at 4 °C until dissection to determine the initial mf intake; henceforth referred to as “Zero hour” mf counts. Mortality was monitored twice daily, and we made special note of accidental deaths and/or mosquitoes that escaped (*n* = 13 of 1,354). All immotile mosquitoes on the bottom of the cage were removed and stored at 4 °C until they were dissected to assess infection status. Accurate parasite counts could not be made in some mosquitoes due to decomposition/drying prior to refrigeration and a missing value was assigned for the parasite loads of such mosquitoes (*n* = 412 of 1345). Dissection and identification of *D. immitis* larval stages were carried out using standard protocols^41^. Mosquitoes in the first and second replicates were censored at 21 and 65 dpf, respectively. The minimum extrinsic incubation period (EIP), the average day at which infective L3 larvae were detected across all populations, was determined to be 13 days (Range: 11-17 dpf; Mean±SD = 12.88±2.10 dpf).

### Statistical analyses

All statistical analyses on were carried out using R 3.3.3 (R Foundation for Statistical Computing). The *A. aegypti* lab strain was used as a “control”, to determine if successful infection, and analyses on *A. albopictus* were carried out after confirming that infection in *A. aegypti* exceeded 85%, as described previously^42^. Results of the regression models (for categorical and continuous variables) were graphed using least-square means and least-square trends, respectively (using the R package LSMEANS^43^). For all analyses the initial model tested expressed *a priori* factors including second order interactions (see specific details below), the best fit model was selected based on lowest AIC while removing statistically non-significant factors, except if the factors were the primary focus of the analysis. ***Exposure to mf:*** We tested for variation in zero-hour mf counts between cages due to the effects of Replicate (i.e. 1 and 2), Dose (i.e. 15 and 30 mf/μL blood), Population, and the interaction between Dose and Population using ANOVA. These results indicated that, within each Dose, parasite exposure differences among populations were negligible (Fig. S2). ***Survival:*** Differences in mortality hazard among the uninfected and infected mosquito lines were considered to be reflective of underlying differences in survival due to unexamined environmental factors (i.e., vigor^44^) and cost of infection, respectively. The Kaplan-Meier method was used to visually compare survival functions for each population at each Dose. Cox Proportional Hazard Mixed Effects models (CMM; implemented in the R package COXME^45,46^) were used to test the effects of Population and Dose and their interactions, on mosquito survival. Replicate was included as a random factor. Escapees and/or accidental deaths were treated as (right) censored data. Detection of significant Population and Population × Dose on mortality hazard was indicated that the populations differed in terms of differences in vigor and infection costs, respectively. ***Resistance:*** Differences in resistance between populations was measured as differences in parasite load at a given time point (when all populations have been initially exposed to the same number of parasites). Since resistance increases the rate at which parasite are killed, lower parasite loads are indicative of greater resistance. We used a Generalized Linear Model (GLMER; as implemented in the R package LME4^47^) with a negative binomial error distribution (and log link) to model the total number of parasites (Parasite Load) as an effect of Population and Day (the day at which the individual died or was censored) and their interactions. All models included the zero hour counts as an offset term, and hence we were able to estimate resistance as the proportional reduction in parasite load at EIP. All models included Replicate as a random factor. A significant effect of Population × Day indicated that the mosquito lines differed in terms of resistance to infection (i.e., the rate at which they killed parasites). ***Tolerance:*** Differences in tolerance between groups of individuals has traditionally been measured as differences in the slopes reflecting a fitness parameter in relation to parasite load. Since the tolerance reduces the negative fitness effects at a given parasite burden, shallower slopes are indicative of greater tolerance^3^. However, unlike microparasites, marcoparasites do not reproduce in the intermediate host (as in the case of *D. immitis* in the vector). Consequently, assuming a constant level of initial parasite exposure, a unit increase in the number of live parasites is necessarily associated with a unit decrease in the number of killed parasites. When populations differ in terms of resistance, ignoring the fitness benefits (or costs) associated with killed parasites will necessarily confound the effects of resistance and tolerance. Thus, we measured the effect of the number of live parasites on population-specific mortality hazards (using CFM) accommodating the number of killed parasites in the model. Variation in this effect (slope and intercept) signified variation among populations in their tolerance. We used parasite counts at the time of mortality or censoring as the measure of the number of live parasites. In case of missing parasite data we interpolated the number of live parasites based on the predictions of the GLMER model (see above). The number of killed parasites at each time point was based on the difference between the number of live parasites and the population-and dose-specific zero-hour counts. A significant effect of Population × Live Parasites (accommodating the effect of resistance) indicated that the mosquito lines differed in terms of tolerance against parasite-mediated pathology. Likewise, significant effect of Population × Killed Parasites was considered to be indicative that the mosquito lines differed in terms of tolerance against immune-mediated pathology. The simultaneous analysis of killed and live parasites, allows us to estimate their unique contributions to survival. ***Vectorial capacity:*** The ability of a mosquito to transmit filarial parasites has traditionally been measured as the proportion of microfilaria ingested that yield infective larvae (i.e., L3 larvae in the head and/or proboscis) amongst mosquitoes surviving the EIP^21^. However, the above measure of vector efficiency ignores mortality through the EIP (i.e., individuals dying prior to the EIP that have no infective larvae^21^). We thus estimated overall vectorial capacity as the joint probability of surviving to EIP and the risk of having L3 in mosquitoes surviving to EIP. This index of vectorial capacity was estimated using zero-inflated negative binomial regression approach. Briefly, the model for vectorial capacity consisted of two submodels: (i) we used a GLMER with binomial error distribution to model the probability of a mosquito surviving to EIP (i.e., non-survivors have no infective parasites) as a function of Dose, Day and Population; (ii) we used a GLMER with negative binomial error to model the risk of infection with an infective L3 larva in the head and/or proboscis of the surviving mosquitoes^21^. To calculate overall vectorial capacity we multiplied the probability of survival to EIP with the relative risk of harboring an L3 larva, and estimated the standard errors (and confidence intervals) of this measure using parametric bootstrap (as implemented in the R package LME4^47^).

### Ethics Statement

All procedures related to vertebrate animals were carried out as approved by NIH Animal Care and Use Committee under protocol LMVR102.

## Acknowledgments

We thank Erica Burkman and Andrew Moorehead (NIH-NIAID Filariasis Research Reagent Resource Center) for their help and advice on *D. immitis*. We are grateful to Dwight Bowman (Cornell University), Melissa Beall and Leif Lorentzen (IDEXX Laboratories) for sharing dog seroprevalence data. We thank Andre Laughinghouse, and Kevin Lee for their help in the insectary. We thank Jeannine Dorothy (Department of Agriculture, MD) for advice and networking with mosquito control authorities. We are indebted to numerous people for help during field sampling, especially: Joseph Marhefka, Evaristo Miqueli (Broward County Mosquito Control, FL), Eric Naguski (Dauphin County Conservation District, PA), Matt Helwig, Michael Hutchinson, Andrew Kyle (Department of Environmental Protection, PA), Michael Cantwell (Department of Agriculture, MD), Jayne Deichmeister, David Gaines (Department of Health, VA), Bob Betts (Escambia County Facilities and Public Works Bureau, FL), MacArthur Dunn (Gadsden County Mosquito Control, FL), Randy Buchanan, Lane Carr (Henrico County Department of Public Works, VA), Gene Lemire (Martin County Mosquito Control District, FL). We thank Peter Armbruster (Georgetown University) and Andrew Read (Pennsylvania State University) for reviewing a draft of the manuscript. For help and support we are also grateful to Phil Lounibos (Florida Medical Entomology Laboratory), Paul Leisnham (University of Maryland), Kevin Caillouet (Virginia Commonwealth University) and colleagues at NIH/NIAID, especially: Sasisekhar Bennuru, Siddhatha Mahanty (Laboratory of Parasitic Diseases), Bob Gwadz, Diana Huestis, Alvaro Molina-Cruz, José Reibeiro (Laboratory of Malaria and Vector Research).

## Authors contributions

Conceptualization, G.D. and T.L.; Methodology, G.D., K.D.W. and T.L.; Investigation, G.D. and K.D.W.; Writing – Original Draft, G.D.; Writing – Review & Editing, G.D., K.D.W. and T.L.; Funding Acquisition, G.D. and T.L.; Supervision, T.L., Competing interests: Authors declare no competing interests.

## Data and materials availability

All data is available in the manuscript, supplementary materials or from the authors.

## List of Supplementary Materials

Figs. S1 to S10

Tables S1 to S6

